# Four-dimensional neural space for moral inference

**DOI:** 10.1101/2025.05.28.656268

**Authors:** Jinglu Chen, Severi Santavirta, Vesa Putkinen, Paulo Sérgio Boggio, Lauri Nummenmaa

## Abstract

Intuitive moral inference enables us to evaluate moral situations and judge their rightness or wrongness. Although Moral Foundations Theory provides a framework for understanding moral inference, its underlying neural basis remains unclear. To capture spontaneous neural activity during moral inference, participants were instructed to watch a film rich in moral content without making explicit judgments while undergoing fMRI scanning. Independent participants evaluated the moment-to-moment presence of twenty moral dimensions in the film. Correlation and consensus cluster analyses revealed four independent main moral dimensions: virtue, vice, hierarchy, and rebellion. While each dimension exhibited unique neural activation patterns, the temporoparietal junction and inferior parietal lobe were activated across all types of moral inference. These findings establish the low-dimensional nature for the neural basis of intuitive moral inference in everyday settings.

## Introduction

Every day, people engage in complex social interactions that require moral evaluation of their own and others’ actions. Should we donate money to charity, should we follow our father’s advice, or should we return the wallet we found on the street to the police station? These decisions and moral judgments are driven by quick, automatic intuitions and emotions (Chen et al., 2024; Haidt, 2001). Because there is no objective right or wrong in the nature, cultures have shaped various sets of moral values for promoting their integrity, and the differences in the moral value codes are primarily products of disparate developmental environments (Haidt et al., 1993; Park et al., 2022; van Hemert et al., 2007). These codes are engaged automatically to evaluate the moral value of others’ actions, leading to emotional responses supporting subsequent affiliative or rejective behavior (Haidt, 2001). Moral reasoning is an intuitive process guided by abstract moral principles and specific life experiences that individuals use to make decisions regarding right and wrong (Ellemers et al., 2019; Haidt & Joseph, 2004). At the individual level, morality promotes personal growth, as those with more advanced moral reasoning are more likely to maintain cooperative behaviors. At the societal level, moral reasoning motivates individuals to advocate for justice and challenge inequities (Killen & Dahl, 2021; Miranda-Rodríguez et al., 2023).

The Moral Foundations Theory (MFT) postulates six core dimensions for moral evaluation, and these universal "intuitive ethics" shape moral judgments across populations (Haidt & Joseph, 2008). These dimensions include care/harm, fairness/ cheating, loyalty/betrayal, authority/subversion, sanctity/degradation, and liberty/oppression. Care vs. harm pertains to actions that nurture and support others’ well-being versus those that cause physical, emotional, or psychological damage. Fairness vs. cheating focuses on evaluating whether actions promote justice, equality, and reciprocity. Loyalty vs. betrayal relates to steadfast commitment and dedication to others versus violating or disregarding established trust and obligations. Authority vs. subversion serves as a foundation for evaluating the moral importance of social hierarchy, strong leadership, and tradition. Sanctity vs. degradation describes the moral weighting of purity, sacredness, and good manners. Liberty vs. Oppression describes the ability to act freely and independently. The relative importance of these moral foundations to individuals or societies influences people’s moral choices. For instance, individuals with differing political ideologies utilize moral foundations in distinct ways when making moral judgments, which can lead to divergent political views and social orientations (Cornwell & Higgins, 2013; Crowson & DeBacker, 2008; Federico et al., 2013; Graham et al., 2009).

Most neuroimaging studies on moral reasoning have however focused on the neural basis of the moral decision-making process, rather than the neural basis of the moral foundations. The prevalent paradigm involves presenting moral dilemmas and evaluating moral violations or moral decision-making (FeldmanHall et al., 2014; Greene et al., 2001, 2004; Han et al., 2014, 2016; Harenski et al., 2008; Hopp et al., 2023; Parkinson et al., 2011; Reniers et al., 2012; Schneider et al., 2013; Shenhav & Greene, 2014). The pivotal fMRI study on the brain basis of moral reasoning demonstrated that emotion plays a significant role in moral judgment. The participants were presented with two types of moral dilemmas—those involving personal emotional engagement and those that did not—and non-moral dilemmas during brain scanning. The results revealed that emotionally charged moral reasoning activated the brain’s emotion circuit more effectively than the other two dilemmas, in the medial frontal gyrus, posterior cingulate gyrus, and angular gyrus. Conversely, when the dilemmas involved less personal emotion, regions associated with working memory, including the middle frontal gyrus and parietal lobe, were more active (Greene et al., 2001).

Subsequent studies have further investigated moral reasoning under similar conditions, revealing the involvement of a distributed network of cortical and subcortical brain regions. The orbitofrontal cortex is involved in theory of mind and reward learning (Greene & Haidt, 2002; Moll et al., 2002; Yoder & Decety, 2018), while the dorsolateral prefrontal cortex is crucial for cognitive control and analyzing moral situations (Greene et al., 2004; Mendez, 2009; Reniers et al., 2012). The anterior cingulate cortex is activated during moral decision making and inference (FeldmanHall et al., 2012; Han et al., 2016). In the temporal lobe, the temporal pole is activated in response to moral transgressions (Finger et al., 2006), and the superior temporal sulcus is important for emotion cognition and theory of mind (Cáceda et al., 2011; Greene et al., 2004; Harenski et al., 2008; Mendez, 2009; Moll et al., 2002; Prehn et al., 2008). The precuneus plays a role in the moral appraisal processes underlying moral reasoning (Harenski et al., 2008; Reniers et al., 2012; Schneider et al., 2013; Young & Saxe, 2008). Additionally, the thalamus is activated by morally unpleasant pictures (Moll et al., 2002), the striatum is involved in moral value evaluation (Shenhav & Greene, 2010; Yoder & Decety, 2018), and the amygdala participates in various aspects of moral reasoning, especially in affective assessment (Berthoz et al., 2006; FeldmanHall et al., 2012; Mendez, 2009; Moll et al., 2002; Schneider et al., 2013; Shenhav & Greene, 2014; Sommer et al., 2010). The insula is engaged in emotion processing and outcome probability assessment (Greene et al., 2004; Mendez, 2009; Shenhav & Greene, 2010), and the temporoparietal junction is crucial for understanding others’ mental states in moral reasoning (FeldmanHall et al., 2014; Finger et al., 2006; Greene et al., 2004; Harenski et al., 2010; Mendez, 2009; Schneider et al., 2013; Sommer et al., 2010; Wasserman et al., 2017; Young & Saxe, 2008).

These fMRI studies demonstrate that neural systems supporting both social cognition and emotions play crucial roles in moral judgments (Greene & Haidt, 2002). However, controlled studies based on artificial moral dilemmas cannot fully capture the natural variation of moral evaluations that occur frequently in everyday life (Wilhelm Hofmann et al., 2015). Because many of the stereotypical situations presented in the moral dilemmas tasks are such that may never be encountered in an average person’s life, researchers have begun to suggest that more ecologically valid approaches should be employed in future research (Graham, 2014).

### The current study

Here, we investigated the neural basis of moral inference during natural vision. Given the inherently spontaneous nature of morality, we departed from traditional self-report paradigms that rely on explicit moral reasoning judgments (Khoudary et al., 2022). Instead, we employed an approach that captured participants’ haemodynamic activation during spontaneous moral inference. Participants viewed a feature film with morally complex and engaging scenes while undergoing functional magnetic resonance imaging (fMRI). The film was dynamically annotated for the moment-to-moment presence of the core moral dimensions postulated in MFT. Modelling the haemodynamic activation parametrically with the time series of the intensities of moral dimensions allowed us to map the brain basis of spontaneous moral processing during complex and contextually rich social scenarios. We further adopted a data-driven approach to examine whether spontaneous moral inference involves six distinct dimensions or whether it is represented by higher-level concepts. The results indicated that moral dimensions in the data can be reduced to four distinct evaluative main dimensions: Virtue, Vice, Hierarchy, and Rebellion. Each of these main dimensions was associated with overlapping, yet distinct neural activation patterns across large-scale cortical and subcortical networks.

## Materials and methods

104 Finnish-speaking volunteers with normal or corrected to normal vision were recruited for the fMRI study. The exclusion criteria were BMI below 20 or above 30, current use of medications affecting the central nervous system, neurological or psychiatric disorders, mood or anxiety disorders, substance abuse, and standard MRI exclusion criteria. Clinically relevant structural brain abnormalities were ruled out by a consultant neuroradiologist based on anatomical MR images. Altogether, eight participants were excluded from the analysis: two due to gradient coil malfunction, two due to abnormalities in structural MRI, and four because of visible motion artefacts in the preprocessed fMRI data. The final sample comprised 96 participants (47 females, mean ± SD age: 31.5±9.36 years, range 20-57). The study was approved by the ethics board of the hospital district of Southwest Finland, and it was conducted according to Good Clinical Practice and the Declaration of Helsinki. An independent sample of 43 healthy Finnish volunteers with normal or corrected-to-normal vision were recruited (35 females, mean ± SD age: 25.3 ± 7.21 years, range 18–53) to evaluate the moment-to-moment presence of moral dimensions from the movie stimuli. All subjects signed a written informed consent and were informed of their right to withdraw at any time without providing a reason. The participants were compensated for their time and travel expenses.

### Stimulus

The Finnish feature film Käsky (*Tears of April* in English, 70min11s) was used as the stimulus since it displays an emotionally intense and morally complex story set during the Finnish Civil War (Louhimies, 2008). This kind of natural audio-visual stimulation has previously been established as a powerful tool for mapping various social and emotional functions (Karjalainen et al., 2017; Lahnakoski et al., 2012; Nummenmaa et al., 2021; Santavirta et al., 2023). In the behavioural experiment, the stimulus was presented in consecutive eight-second-long chunks while the neuroimaging data was acquired in two sessions (32 min 44 s and 37 min 27 s) with a short break in between.

### Evaluation of moral and emotional dimensions

The evaluated moral dimensions: Care/Harm, Fairness/Cheating, Loyalty/Betrayal, Liberty/Oppression, Authority/Subversion, and Sanctity/Degradation were based on the MFT (Haidt & Joseph, 2008; Iyer et al., 2012). We also incorporated additional dimensions, including Individualism (harm versus benefit to the individual) and Collectivity (harm versus benefit to the group), which serve as overarching features of moral reasoning (Graham et al., 2009). Because moral inference is influenced by basic emotions, we included dimensions for pleasure, displeasure, arousal, and calmness, which capture a substantial proportion of variance in basic emotion ratings (Russell, 1980).

Furthermore, we included simple dimensions of “moral righteousness” and “moral injustice” as umbrella terms to capture the rudimentary “gut feeling” of moral evaluations. In total, we collected data for 20 dimensions (**Table 1**).

**Table 1.**
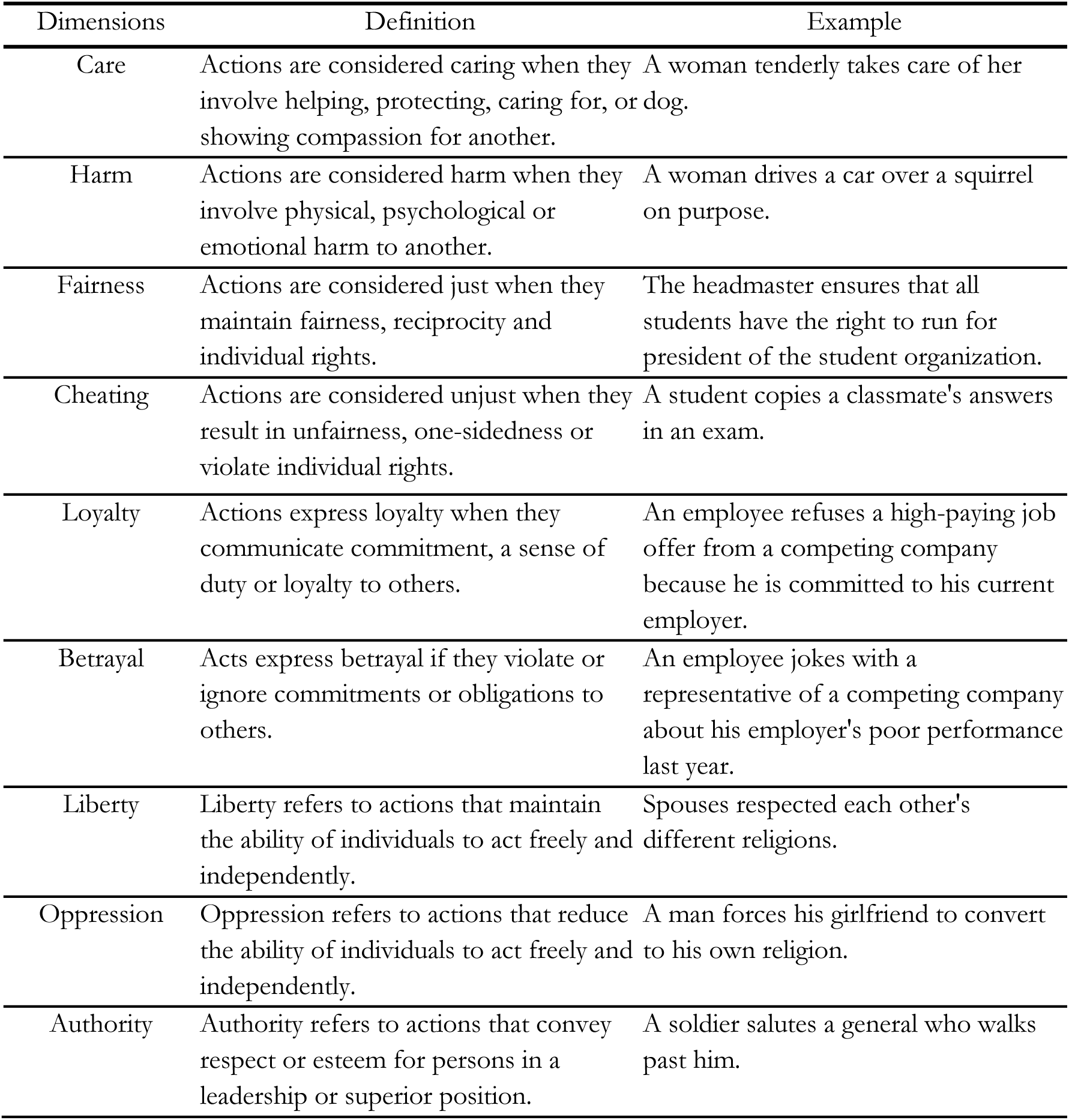
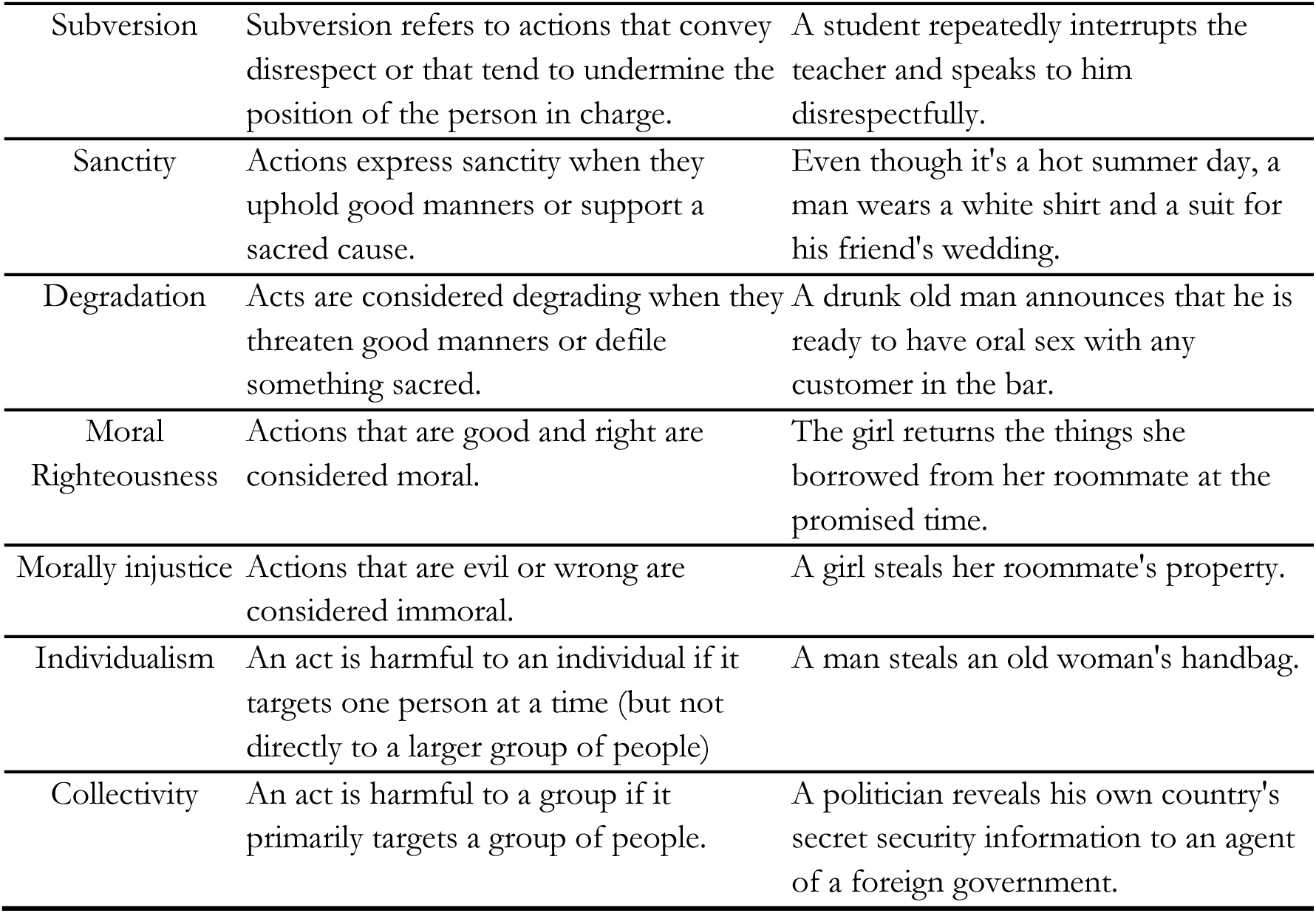
Moral dimensions, as well as their definitions and examples.

In the behavioral experiment, participants were instructed as follows: Your task is to assess whether the film depicts aspects related to the dimensions important for moral evaluation. The definitions and examples for each dimension were also presented to the participants. The movie was split into eight-second clips. Two interleaved splits were created with 4-second difference in starting times, thus yielding 4-second temporal resolution when the rating data were combined across the splits. These splits were presented to different participants to provide mean moral evaluations of the whole movie at a four-second temporal resolution (**Figure S1**). Each participant was randomly assigned to evaluate one of the two sets of interleaved movie clips for 10 of the 20 evaluated dimensions. After viewing a movie segment, participants rated the presence of elements related to the predefined dimensions on a scale from 0 (not at all) to 10 (very much) (See **Figure S3** for the ratings corresponding to the 20 dimensions). The participants evaluated the moral contents of the movie in the laboratory and the ratings were collected using the Gorilla platform (https://gorilla.sc/). The ratings from all participants were then averaged to produce the final moral inference scores. See **Figure 1** for the data collection and analysis procedures (for more details, see section *Behavioral data collection and analysis* in the supplementary materials).

**Figure 1.**
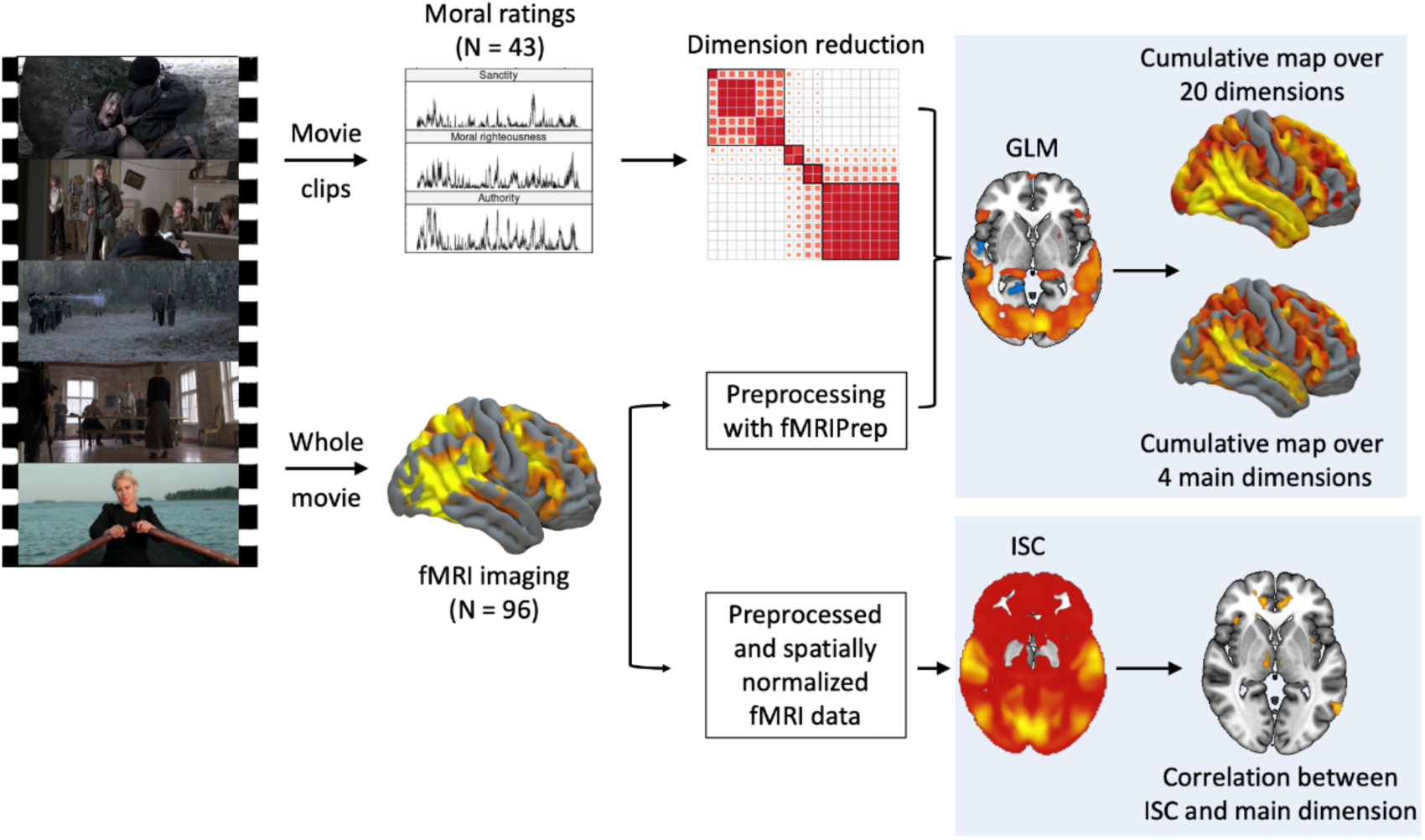
Data acquisition and processing.

### Dimension reduction analysis for the moral dimensions

To determine whether the theoretical dimensions of moral inference form independent evaluative dimensions or larger evaluative categories in the movie stimuli, we conducted a consensus clustering analysis. First, a correlation matrix of the mean time series of 20 moral evaluative dimensions was calculated. Temporal correlation quantifies the co-occurrence of dimensions in the movie, and high correlations would suggest shared or overlapping processes when making the evaluations. Consensus clustering analysis was then conducted on the correlation matrix using the DiceR package (Chiu & Talhouk, 2018). Consensus approach ensured stable clustering structure by running 1,000 independent iterations with unique subsamples of the data and over different numbers of clusters (from 2 to 8 clusters).

### fMRI acquisition and preprocessing

During the fMRI experiment, the participants were instructed to stay still and watch the movie attentively. The experiment was run using Presentation software (https://www.neurobs.com/). Visual stimuli were presented with NordicNeuroLab VisualSystem binocular display. Sound was conveyed with Sensimetrics S14 insert earphones. Before the functional run, sound intensity was adjusted for each subject so that it could be heard over the gradient noise.

Functional MRI data were acquired with the Phillips Ingenuity TF PET/MR 3-T whole-body scanner at the Turku PET Centre. T1-weighted high-resolution structural images (1mm^3^ resolution, TR = 9.8 ms, TE = 4.6 ms, flip angle = 7°, FOV = 250 mm, 256 × 256 reconstruction matrix) and T2*-weighted functional images (TR = 2600 ms, TE = 30 ms, flip angle = 75°, FOV = 240 mm, 80 × 80 reconstruction matrix, 62.5 kHz bandwidth in EPT direction, 3.0 mm slice thickness, 45 interleaved axial slices acquired in ascending order without gaps) were collected and preprocessed with fMRIPrep 1.3.0.post2 (Esteban et al., 2019).

Preprocessing of the T1-weighted images included correction for intensity non-uniformity (INU), distributed with ANTs 2.2.0 and used as a reference later. The images were skull-stripped and reconstructed using recon-all for brain surfaces. The brain mask was refined with a custom variation of the method to reconcile ANTs-derived segmentations of the cortical gray matter of Mindboggle. Spatial normalization to the ICBM 152 Nonlinear Asymmetrical template version 2009c and brain tissue segmentation were also performed. The following preprocessing was performed on the functional data: slice-time correction was done using 3dTshift, and the data were resampled to fsaverage5. They were then resampled to their native space for head-motion and susceptibility distortions. Spatial smoothing was applied with an isotropic Gaussian kernel of 6mm FWHM. Automatic removal of motion artifacts was performed using ICA-AROMA.

### General linear model analysis for the fMRI

FMRI data were analyzed in SPM12 (Wellcome Trust Center for Imaging, London, UK, https://www.fil.ion.ucl.ac.uk/spm/). To map the regions associated with moral inference, we first standardized (0 mean, 1 SD) the evaluative rating time series. Next, we averaged the ratings of moral dimensions within each cluster identified in the dimension reduction analysis to get a cluster-level time series of the prevalence of major moral events in the films. These time series were convolved with canonical HRF and re-standardized before incorporating them into first-level general linear models (GLM).

Additional details on the variance inflation factors (VIF) between the convolved cluster time series can be found in the supplementary materials (*Correlations and VIF values for the movie*). In the full-volume analysis, contrast images were first generated separately for each participant to examine the main effects of the four clusters for moral inference.

These contrast images were then subjected to a second-level analysis (one-sample t-test) to estimate the population-level effects. Finally, the voxel-wise results were corrected for multiple comparisons using family-wise error (FWE) at the cluster level. Specifically, statistical maps were thresholded at *p_unc_* < 0.001, identifying the FWEc minimum cluster-size value for FWE correction at the cluster-size level, and then thresholded again at *p_unc_* < 0.001 and k = FWEc.

In the ROI analysis, 56 bilateral anatomical ROIs were extracted from the AAL2 atlas (Rolls et al., 2015). A one-sample t-test on the mean ß-weights of each ROI was used to assess statistical inference on the group-level. ROI-analysis results were considered significant with p < 0.05 (Bonferroni-corrected for multiple comparisons).

### Cumulative maps for moral inference

To identify brain regions activated during different types of moral inference, we summarized the results by calculating a cumulative map (binarized sum of significant activations) over all 20 original moral dimensions and four main dimensions separately to reveal a fine-grained gradient in the brain basis of moral inference. Modelling and statistical thresholding for individual moral dimensions were conducted similarly to the main analysis. Subsequently, two cumulative maps of the binarized BOLD beta maps were computed for original dimensions and four main dimensions, highlighting regions with the most pronounced BOLD responses associated with moral inference.

Moral inference occurs in social situations and is inherently based on socioemotional factors. To identify how the brain responses to moral inference overlap with general social perception networks, we conducted similar cumulative analysis for 44 consistently evaluated social perceptual features that were previously evaluated for the same stimulus (Santavirta et al., 2023). Subsequently, we utilized 56 bilateral anatomical regions of interest (ROIs) from the AAL2 atlas to calculate the average cumulative proportion within each ROI for two moral inference cumulative maps and social perception map. Finally, we computed correlations at the ROI level between the social perception cumulative map and the cumulative maps of two moral inference dimensions.

Finally, we examined whether moral inference synchronizes neural activations across participants. We conducted a supplementary time-window intersubject correlation analysis (ISC) and modelled the time-varying neural synchronization with the four dimensions for moral inference (see supplementary materials: *Inter-subject correlation analysis* for more detail).

## Results

### Main dimensions of moral evaluation

Consensus clustering analysis identified a four-cluster structure that was both theoretically meaningful and supported by the data-driven clustering for the 20 evaluated moral dimensions. These four clusters were chosen for further analysis of the fMRI data (**Figure 2**). They were labeled as labeled as ’**virtue**’ (sanctity, care, loyalty, fairness, moral righteousness, liberty, pleasure, calmness), ’**hierarchy**’ (authority, collectivity), ’**rebellion**’ (subversion, individualism), and ’**vice**’ (betrayal, arousal, degradation, oppression, displeasure, moral injustice, harm, cheating). The cluster-level time-series are shown in **Figure 3**. The correlations between the resulting four cluster time series ranged from - 0.34 to 0.54, and the VIF values ranged from 1.16 to 2.40. This shows that the regression coefficients for the four main dimensions are stable in the linear model estimations, and that they can be included in the same regression model given their independence.

**Figure 2.**
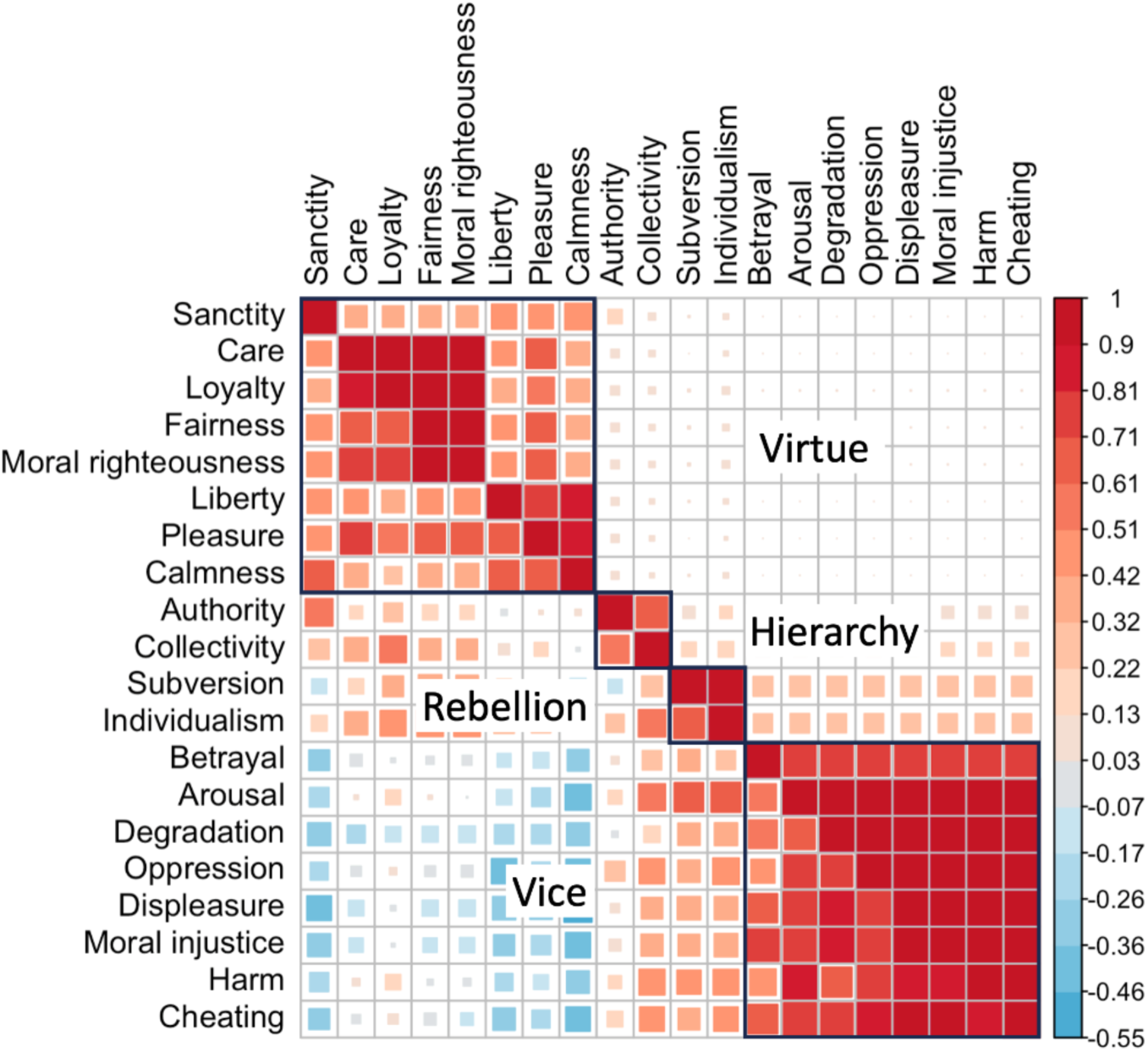
Correlation (lower left) and consensus (upper right) matrix.

**Figure 3.**
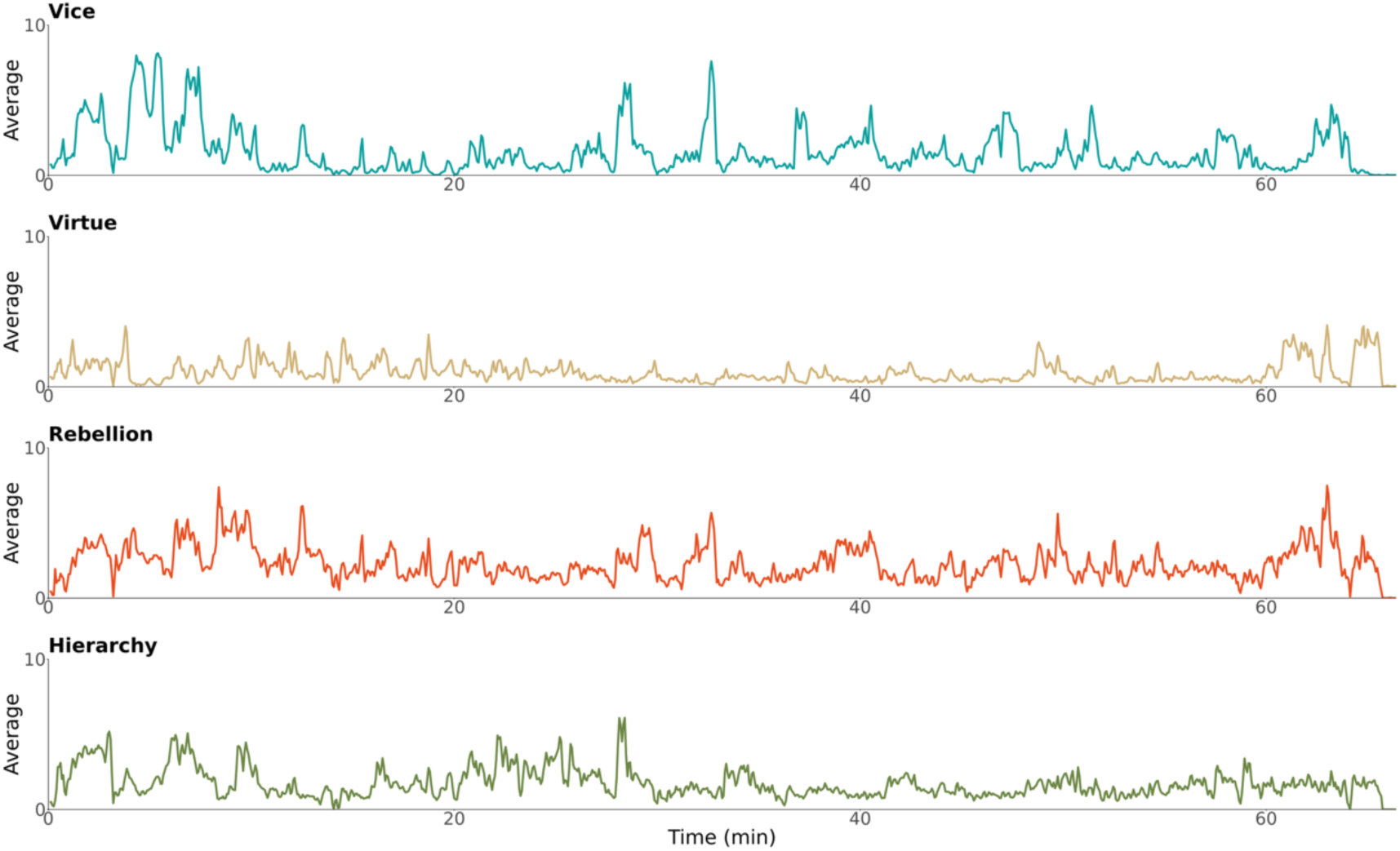
Temporal distribution of mean ratings for four main dimensions throughout the movie.

### Cerebral topography of moral inference

The whole-brain general linear model (GLM) revealed distinct neural activation patterns across four main moral dimensions (**Figure 4**). The **vice** dimension engaged large-scale cortical and subcortical networks, including the medial frontal gyrus, anterior cingulate cortex, precuneus, posterior cingulate cortex, dorsolateral prefrontal cortex, middle frontal gyrus, temporoparietal junction, superior temporal gyrus, orbitofrontal cortex, insula, amygdala, caudata, putamen, thalamus, supramarginal gyrus and angular gyrus. In contrast, the neural activation patterns for **virtue** spanned a smaller area compared to the vice cluster. However, temporal regions, including the superior temporal sulcus and the temporal pole, were responsive to the virtue dimension rather than to the vice dimension. The **rebellion** dimension was associated with both positive and negative neural activations. Positive activation was observed in the medial frontal gyrus, precuneus, posterior cingulate cortex, dorsolateral prefrontal cortex, temporoparietal junction, superior temporal sulcus, superior temporal gyrus, temporal pole, caudate, thalamus, supramarginal gyrus, and angular gyrus. Conversely, the anterior cingulate cortex exhibited negative activation. Finally, the **hierarchy** dimension differed markedly from the other main dimensions, showing primarily negative associations with neural responses. Only few regions, including the amygdala, temporoparietal junction, superior temporal sulcus, superior temporal gyrus, supramarginal gyrus, and angular gyrus showed positive activation.

**Figure 4.**
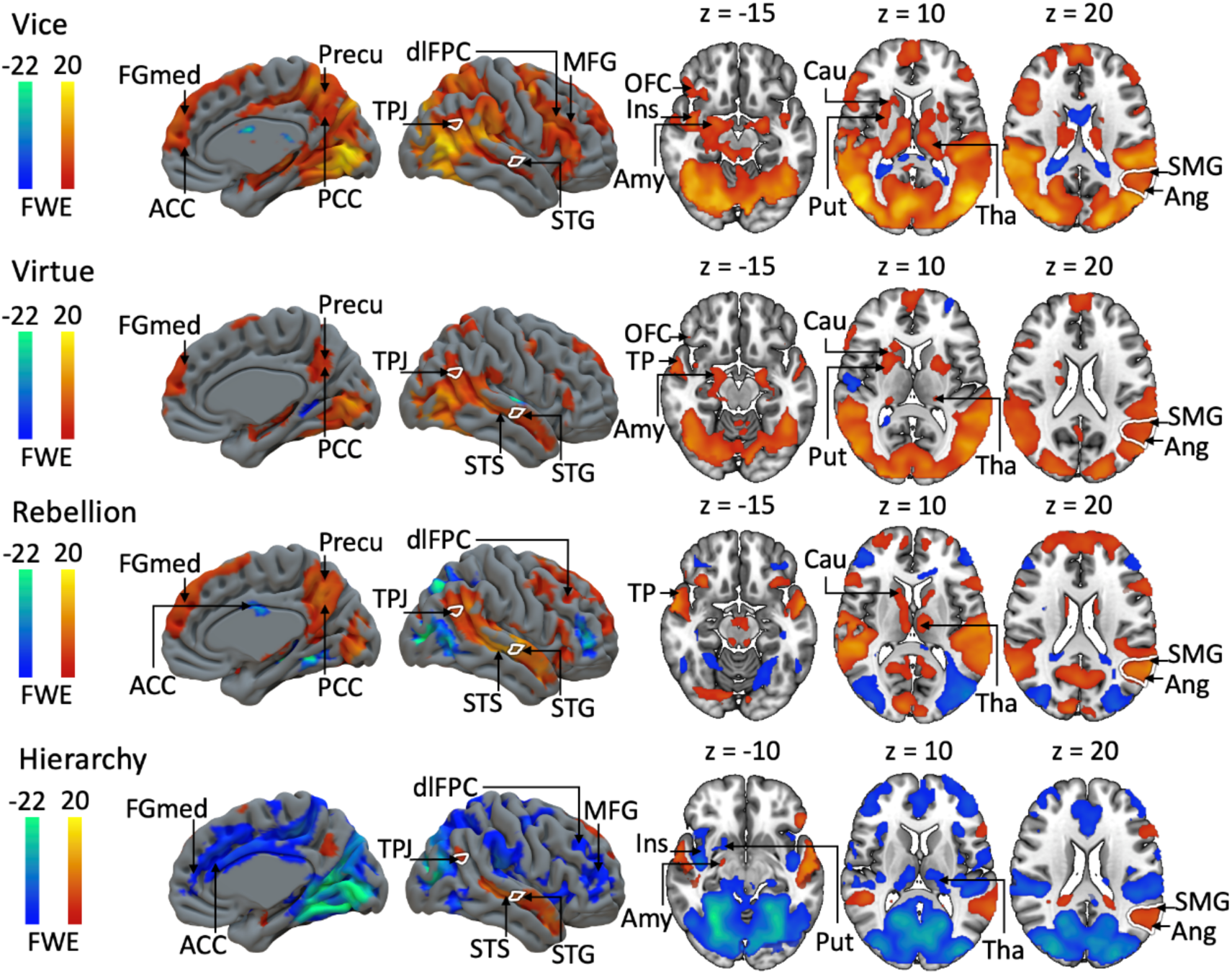
Neural activation patterns associated with the four main dimensions of moral inference (Cluster FWE-corrected at p < 0.05. Initial cluster-forming threshold p < 0.001). Color bars indicate the t-statistic range. FGmed = medial frontal gyrus, Precu = precuneus, ACC = anterior cingulate cortex, PCC = posterior cingulate cortex, dlPFC = dorsolateral prefrontal cortex, MFG = middle frontal gyrus, TPJ = temporoparietal junction, STS = superior temporal sulcus, STG = superior temporal gyrus, OFC = orbitofrontal cortex, Ins = insula, Amy = amygdala, TP = temporal pole, Put = putamen, Cau = caudate, Tha = thalamus, SMG = supramarginal gyrus, Ang = angular gyrus.

Results from the ROI analysis supported the full-volume analysis results (**Figure 5**). Overall, the regional effects were mostly positive for vice, virtue, and rebellion, and negative for hierarchy, except for temporal positive associations. Spatially, the associations between moral inference and haemodynamic activity were most consistently observed in the temporal lobe. Frontoparietal activations were most prominent for vice and rebellion dimensions, while occipital responses were consistently positive for vice and virtue, negative for hierarchy, and mostly absent for rebellion. Finally, subcortical activity was most consistently associated with vice.

**Figure 5.**
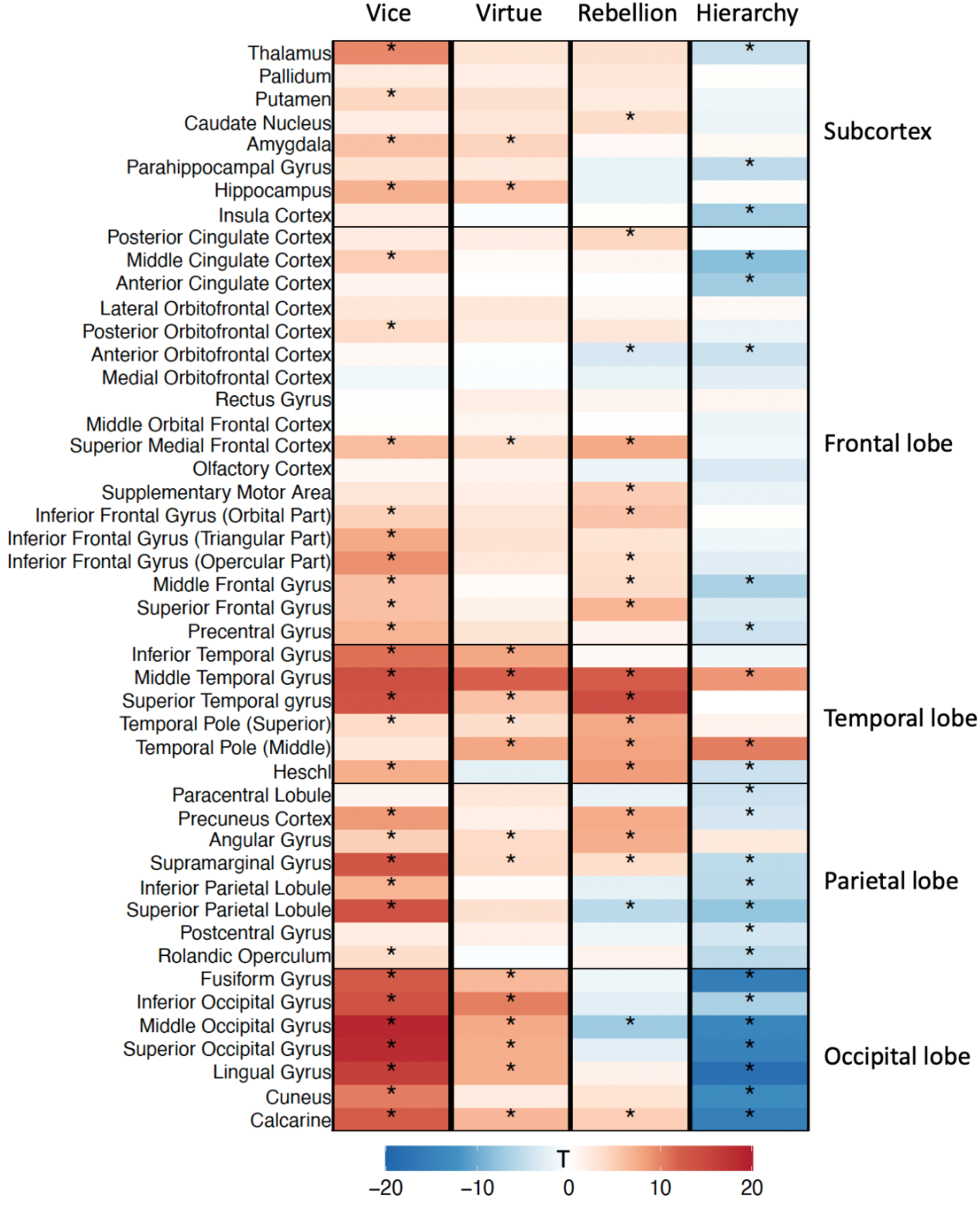
The heatmap displays the t values for regression coefficients corresponding to each ROI and moral inference cluster. Statistically significant ROIs (p < 0.05, Bonferroni corrected independently for each moral cluster) are indicated with an asterisk.

### Domain-general responses during moral inference

The cumulative analysis of all 20 original moral dimensions and the four main dimensions revealed extensive subcortical-cortical activation patterns. The occipital lobe and subcortical regions demonstrated the most consistent activations across both calculation methods. For comparison, the cumulative map of social perception also exhibited widespread activation patterns (**Figure 6**).

**Figure 6.**
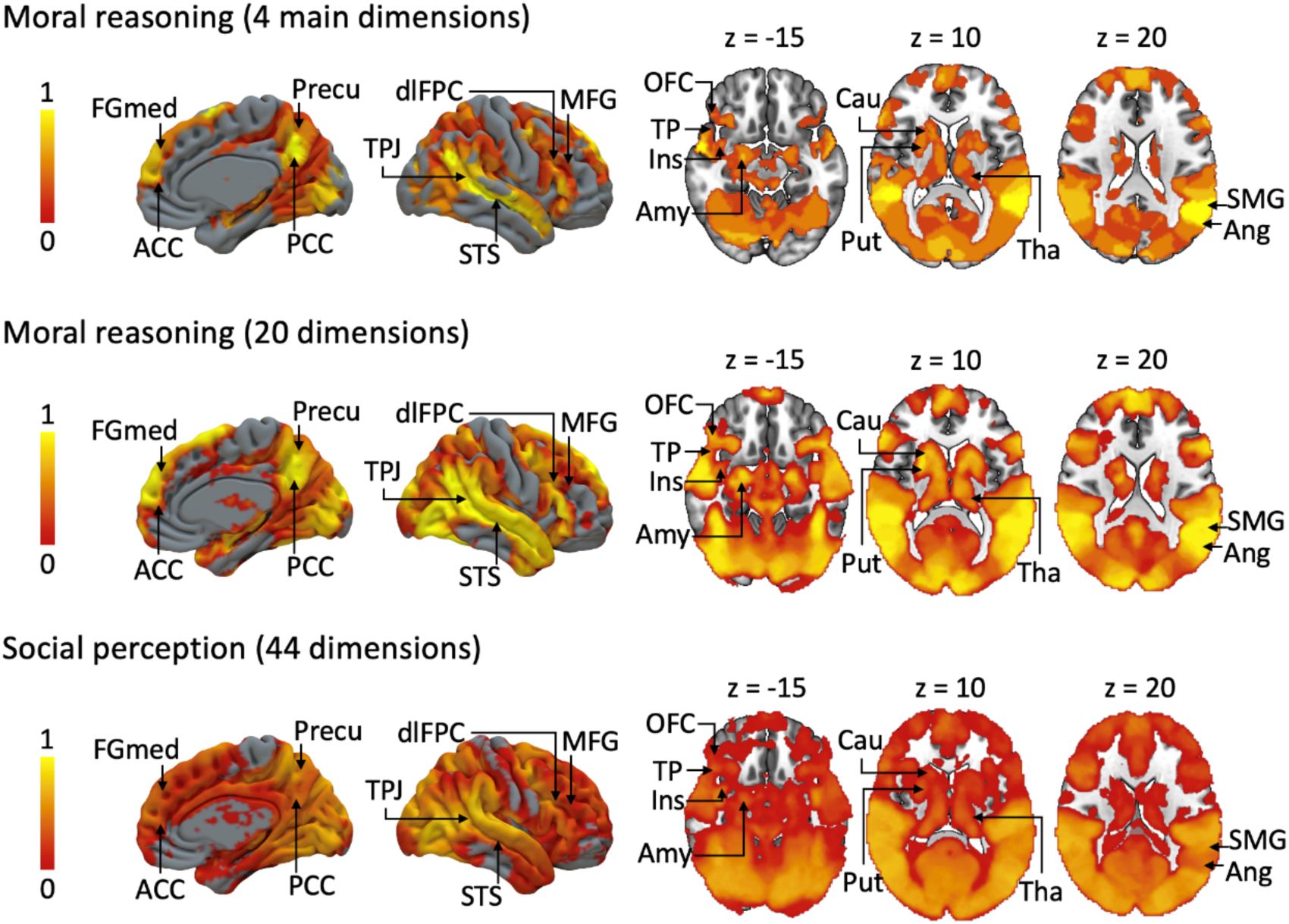
Cumulative activations for moral inference and social perception. The colorbar represents the proportion of dimensions that associated significantly (Cluster FWE-corrected at p < 0.05. Initial cluster-forming threshold p < 0.001) with the neural responses. FGmed = medial frontal gyrus, Precu = precuneus, ACC = anterior cingulate cortex, PCC = posterior cingulate cortex, dlPFC = dorsolateral prefrontal cortex, MFG = middle frontal gyrus, TPJ = temporoparietal junction, STS = superior temporal sulcus, OFC = orbitofrontal cortex, TP = temporal pole, Ins = insula, Amy = amygdala, Cau = caudate, Put = putamen, Tha = thalamus, SMG = supramarginal gyrus, Ang = angular gyrus.

The correlation analysis between moral inference and social perception revealed substantial overlap in activation patterns between these two processes. The temporal, occipital, and parietal lobes demonstrated the most consistent activation across both moral inference and social perception during movie viewing. Notably, subcortical regions exhibited more consistent activation in moral inference (**Figure 7**).

**Figure 7.**
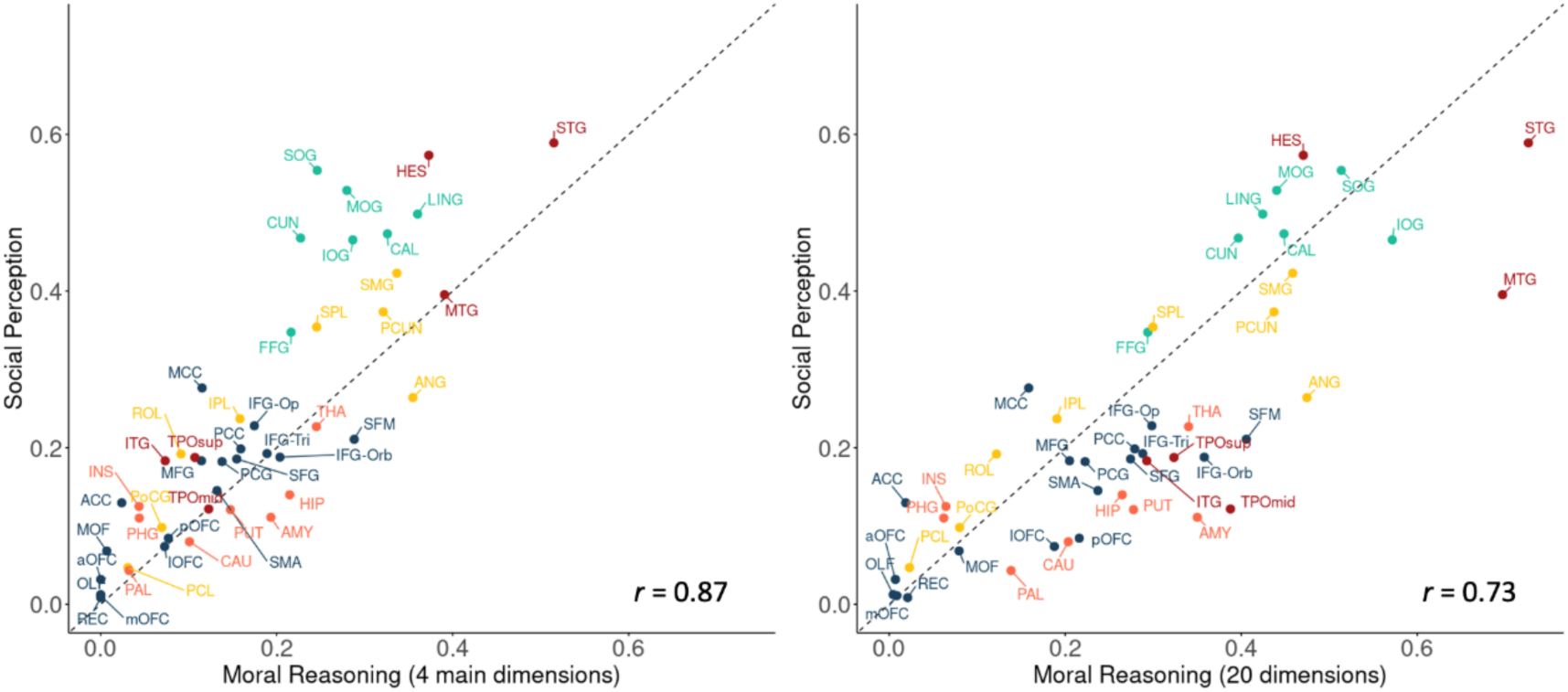
Scatterplot showing the relationship between the proportion of cumulative regional activations for moral inference and social perception. The left panel depicts the correlation using the cumulative map derived from the four main dimensions of moral inference, while the right panel shows the correlation based on all 20 dimensions. Each point represents an anatomical ROI (see **Table S3** for the full name of the regions). The position in relation to the diagonal indicates whether a ROI is more tuned to moral inference (below diagonal) or social perception (above diagonal).

## Discussion

Our main finding was that moral inference during natural vision is based on four main dimensions which have both shared and independent neural bases. Moral inference is often a spontaneous and intuitive process that occurs without conscious deliberation (Haidt, 2001), yet many studies require participants to make decisions regarding moral dilemmas, which may not accurately reflect the underlying moral inference process (Graham et al., 2009; Greene et al., 2001; Hopp et al., 2023; Khoudary et al., 2022). Here, we presented participants with a full-length feature film that was previously annotated for the presence of different moral dimensions. This approach identified four main dimensions of moral inference: Virtue, Vice, Hierarchy, and Rebellion. The participants viewed the movie during fMRI without an explicit task, allowing us to quantify the brain basis of spontaneous moral inference in a naturalistic context. Although the brain activation patterns associated with the four main dimensions showed marked differences, they also shared common neural bases, particularly in the TPJ and inferior parietal lobe. These data highlight the low-dimensional nature of moral inference, with distinct but partially overlapping brain basis across dimensions.

### Four foundations of moral inference

Moral Foundations Theory posits six psychological “systems” that individuals utilize to guide their moral judgments (Haidt & Joseph, 2008; Iyer et al., 2012). To test whether these dimensions are independently evaluated in the naturalistic movie stimulus, we first conducted correlation and cluster analyses on the behavioural moral ratings. This revealed that many of the proposed dimensions are interrelated, suggesting shared or overlapping evaluative processes. Consensus clustering revealed four distinct clusters (**Figure 2**). Sanctity, care, loyalty, fairness, and liberty, from MFT, as well as moral righteousness, pleasure, and calmness, were clustered together. These dimensions indicate favourable or desired moral evaluations, and we thus labelled the cluster as "**Virtue”.** Authority and collectivity were included in another cluster. Authority refers to actions that demonstrate respect or esteem for individuals in leadership or superior positions, while collectivity refers to actions that may impact the well-being of the group rather than just a single individual. These dimensions primarily emphasize the importance of respecting authority and prioritizing collective interests. Consequently, we designated this cluster as "**Hierarchy**," reflecting its focus on behaviours that promote the collective welfare and social hierarchy.

The third cluster encompassed the dimensions of subversion and individualism. Subversion refers to actions that express disrespect or undermine the authority of individuals in charge. The dimension of individualism pertains to actions that are detrimental to an individual. Together, these dimensions highlight behaviours that are either harmful or disrespectful toward individuals. Therefore, we designated this cluster as "**Rebellion**," reflecting its focus on behaviours that influence primarily the individual rather than the group. The last cluster includes the dimensions of betrayal, degradation, oppression, harm, and cheating, as identified in MFT, along with arousal, displeasure, and moral injustice. In contrast to the virtue cluster, the dimensions within this cluster were associated with negative or unfavourable moral evaluations. We labelled this cluster as "**Vice**," reflecting the negative behaviours identified in moral inference. The variance inflation factor values for these four clusters indicated that they were independent from one another. In contrast to the original Moral Foundations Theory, we dismantled the paired dimensions and considered them as independent constructs that might have involved different evaluative processes.

Virtue and Vice reflected how individuals perceived behaviours as either good or bad, morally righteous or wrong. Reasoning about right and wrong is a fundamental aspect of ethical decision making, which serves as a fundamental feature of human morality and plays a pivotal role in maintaining social order (Ellemers et al., 2019). The dimensions of Virtue and Vice represent possible distinct pathways through which individuals process morally right and wrong information, highlighting the complexity of moral cognition. As social animals, humans naturally engage in cooperation, leading to the emergence of groups (Tomasello, 2014). Given the importance of group dynamics to each individual, it is reasonable to conclude that people were particularly sensitive to behaviours that pertained to themselves or their groups. The Hierarchy and Rebellion main dimensions suggested that individuals might have perceived behaviours differently depending on whether they targeted an individual or a group.

### Neural basis of automatic moral inference

Previous imaging research on brain basis of moral reasoning has employed controlled paradigms where participants make deliberate moral judgments in artificial scenarios pertaining specific moral concerns (FeldmanHall et al., 2014; Greene et al., 2001, 2004; Han et al., 2014, 2016; Harenski et al., 2008; Hopp et al., 2023; Khoudary et al., 2022; Parkinson et al., 2011; Reniers et al., 2012; Schneider et al., 2013; Shenhav & Greene, 2014). In contrast, we investigated the automatic, intuitive processes involved in moral inference using a naturalistic movie viewing where participants were not required to provide explicit answers to moral-related questions during brain imaging. Instead, the potential moral questions emerged naturally from the flow of the action in the cinema (Karjalainen et al., 2017; Lahnakoski et al., 2012; Nummenmaa et al., 2021; Santavirta et al., 2023).

The analysis of both the initial 20 dimensions and 4 higher-order main dimensions revealed a widely distributed network of cortical and subcortical regions involved in encoding the moral behaviours depicted in the movie (**Figure 6**). Consistent activation was observed in the medial frontal gyrus, posterior cingulate cortex, temporoparietal junction, superior temporal sulcus, and amygdala – regions previously identified as key nodes of emotion processing in moral reasoning (FeldmanHall et al., 2012; Greene et al., 2001, 2004; Harenski et al., 2008; Mendez, 2009; Moll et al., 2002; Wasserman et al., 2017). Additionally, the dorsal striatum and inferior parietal lobe exhibited consistent activation, aligning with their established roles in working memory and value evaluation in prior studies (Greene et al., 2001; Han et al., 2016; Mendez, 2009; Reniers et al., 2012; Shenhav & Greene, 2010). These activation patterns resemble the brain’s social perception network, and comparison of the presently observed moral judgment networks with previously established network for social perception revealed clear correspondence, as illustrated in **Figure 7**. The more social features a region was responsive to, the more moral dimensions it also responded to. However, there was a clear cortical-subcortical gradient in general-purpose social versus moral processing, in that subcortical regions involved in emotions and motivation exhibited wider tuning profiles for moral versus social perception, highlighting the affective nature of moral information processing and decision making.

The GLM analysis showed that the four main dimensions of moral inference were found to share a common cerebral basis while also exhibiting unique activation patterns.

Consistent activation across all dimensions was observed in the temporoparietal junction (TPJ), superior temporal gyrus, supramarginal gyrus, and angular gyrus across these dimensions. The TPJ, a key region associated with Theory of Mind and understanding others’ mental states (Saxe & Kanwisher, 2003), showed consistent activation in our study, suggesting its role in encoding and integrating mental state information as a foundational process for further moral judgments (Wasserman et al., 2017; Young & Saxe, 2008). The supramarginal gyrus and angular gyrus, both components of the inferior parietal lobe (IPL), have been shown to be active during moral decision-making tasks (Greene & Haidt, 2002; Han et al., 2016; Mendez, 2009; Reniers et al., 2012). The IPL is recognized as a critical hub underlying diverse basic and social cognitive processes. The consistent activation of the IPL across the four dimensions of moral inference suggests a unified function in processing cognitive and social information essential for moral inference. Although the superior temporal gyrus was not frequently reported in previous studies, it has consistently been shown to activate during social perception (Santavirta et al., 2023). Compared to prior controlled experiments, the naturalistic paradigm used in the present study may have more effectively elicited activation of the superior temporal gyrus.

There were also notable differences in the activation patterns across the dimensions. The Vice main dimension showed the most pronounced positive activation, particularly in subcortical regions like the thalamus, striatum, and amygdala (**Figures 4 and 5**). The widespread brain activation observed in response to the Vice main dimension suggests a complex interplay between cognitive and emotional processes in the brain’s reaction to morally questionable or potentially harmful stimuli. Voxel-wise analysis revealed selective activation of the anterior cingulate cortex (ACC), insula, and middle frontal gyrus in response to vice. The ACC, known for its role in cognitive control (Botvinick et al., 2001), demonstrated positive activation, which may indicate that cognitive control systems are particularly engaged when behaviours are perceived as morally wrong or conflict with internal values. The middle frontal gyrus, previously implicated in working memory during moral reasoning tasks (Greene et al., 2001), likely supports this process by maintaining relevant information for moral evaluation. The insula, a critical hub for integrating cognitive and emotional information (Namkung et al., 2017), showed activation that underscores its integrative role in processing information during the detection of moral violations. Additionally, the insula and anterior cingulate cortex-key nodes of the salience network-were observed (Menon & Uddin, 2010). This activation may indicate that situations involving moral violations are particularly salient, requiring the integration of complex information and heightened attention.

Fewer brain regions were activated in response to Virtue compared to Vice, particularly in subcortical areas, the frontal lobe, and the parietal lobe. This pattern of reduced activation may suggest that the detection of morally right behaviour requires less cognitive and emotional effort compared to the processing of morally questionable or harmful actions. The caudate nucleus and posterior cingulate cortex were activated consistently in response to the Rebellion in both analyses, but not for other main dimensions (**Figure 4** & **Figure 5**). Previous research has shown that these areas exhibit greater activation for personal moral judgments compared to impersonal moral judgments (Greene et al., 2004). In this study, brain activation was notably higher when the film depicted behaviours that caused more harm to individuals. These results suggested that the caudate nucleus and posterior cingulate cortex are involved in detecting immoral behaviours directed at individuals. Responses to the hierarchy dimension exhibited extensive negative activation (**Figure 4** & **Figure 5**). The stimuli, drawn from a war movie featuring disturbing scenes such as shootings, likely evoked feelings of fear among participants. Notably, these disturbing scenarios predominantly involved groups of people. Based on participant ratings, these parts of the movie received relatively higher scores in the authority and collectivity dimensions compared to other segments. The amygdala, which is known to be activated in response to fear-related emotions (Ressler, 2010), showed increased activation when greater harm was inflicted on the group. In contrast, most other brain regions exhibited deactivation. This pattern may suggest that participants experienced a fear response, leading them to avoid processing behaviours that caused harm to the group.

### Limitations

We deliberately wanted to study implicit moral judgments, so a separate sample participated in the rating and fMRI experiment. We do not thus know how well the participant-level moral processing matches with the population level average derived from the fMRI experiment. Despite this, the brain responses to the different dimensions were consistent across participants, indicative of convergence. Finally, although we acquired data from well over one hour of morally evocative scenarios, it is possible that all possible moral dimensions were not comparably present in the stimulus. However, the time series of the main moral dimensions revealed significant variation in all original as well as higher-order dimensions.

## Conclusion

We conclude that a low-dimensional space effectively captures the subjective and neural-level variation in moral inference. Contrary to the Moral Foundations Theory, our findings revealed four distinct foundations that appear to process in parallel within the human brain: Virtue, Vice, Hierarchy, and Rebellion. While each of these main dimensions demonstrated unique neural signatures, they also shared common neural substrates in the TPJ and inferior parietal lobe. Our data suggest that automatic moral inference involves the detection of morally right and wrong information, as well as behaviours directed toward individuals or groups. These findings establish the low-dimensional nature for the neural basis of intuitive moral inference in everyday settings.

## Acknowledgements

This work was supported by Aatos Erkon Säätiö, China Scholarship Council (202106040042), Alfred Kordelin Foundation, and Research Council of Finland (grant #350416).

## Supplementary materials

### Behavioral data collection and analysis

To get compact moral emotion intensity fluctuation during the whole movie, and decrease the fragmenting feeling of watching movie, we created two sets of movie clips. The first set began at the start of the film, with the first clip lasting 4 seconds and all subsequent clips lasting 8 seconds. The second set also started at the beginning but consisted entirely of 8-second clips. As shown in **Figure S1**, groups 1 and 2 viewed the first set of clips, while groups 3 and 4 viewed the second set. There are 10 raters in group1, 12 raters in group 2, 10 raters in group3, and 11 raters in group4.

**Figure S1.**
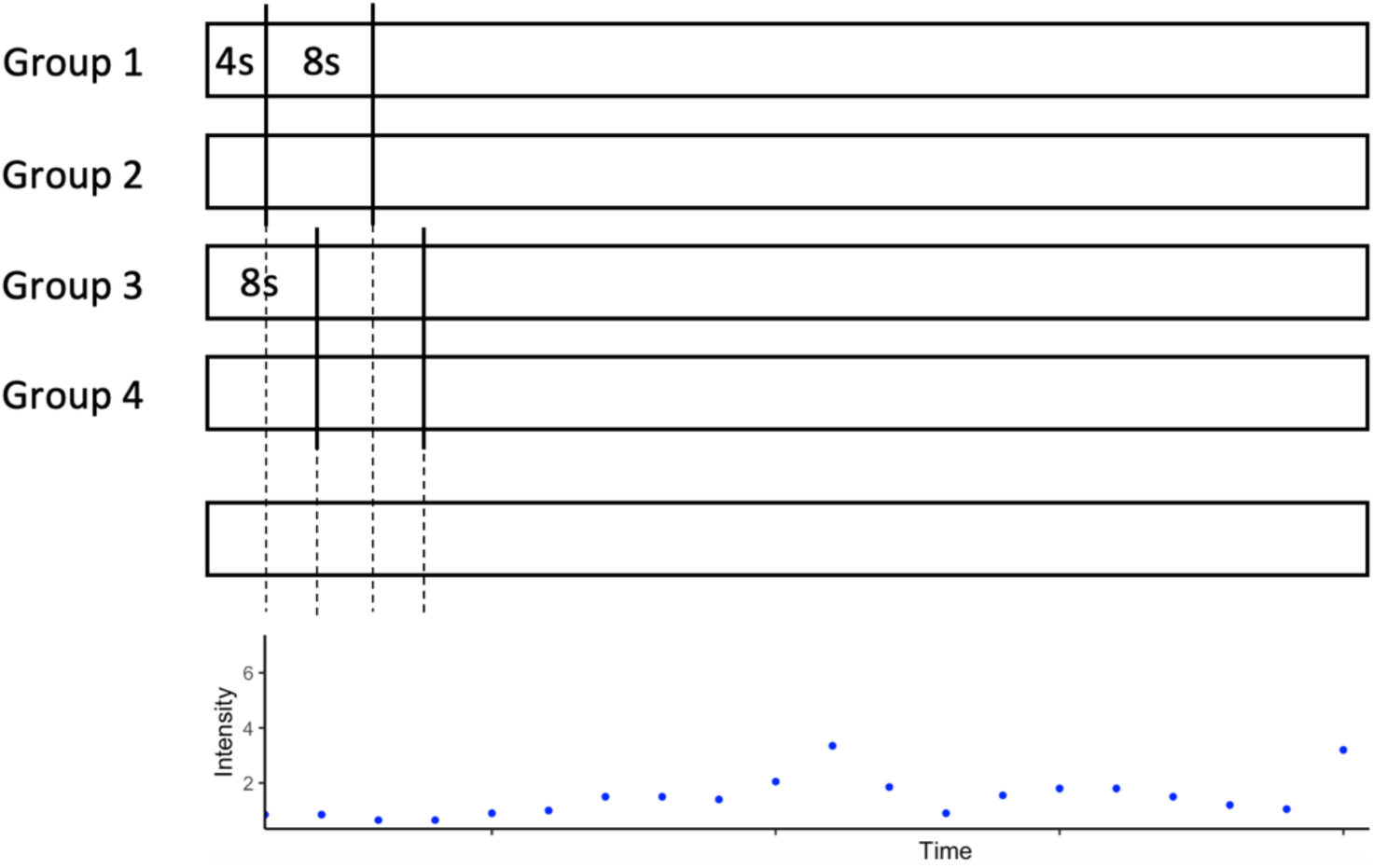
Videos clips viewing in the behavioural experiment. In the behavioral experiment, video clips were segmented differently for participant groups. Groups 1 and 2 initially viewed a 4-second clip, followed by 8-second clips for the remainder of the experiment. In contrast, Groups 3 and 4 consistently viewed 8-second clips throughout. Groups 1 and 3 rated the same set of 10 moral inference dimensions, while Groups 2 and 4 rated the remaining 10 dimensions. Final ratings for each dimension were calculated as the average of all participant ratings. By combining ratings across different time points, we generated ratings for each 4-second time window across all 20 dimensions.

The 20 dimensions of moral inference were divided into two subsets, with participants in each group rating only 10 dimensions per set. In **Figure S1**, groups 1 and 3 rated the first set of moral inference dimensions, while groups 2 and 4 rated the second set. Refer to **Figure S2** for the intraclass correlation coefficients (ICC) within each rating group.

To achieve more accurate estimates of moral emotion intensity that closely reflect participants’ perceptions, we calculated the mean ratings from both relevant groups. For instance, for the first 12 seconds of the film, groups 1 and 3 provided three data points for emotion intensity across 10 dimensions, while groups 2 and 4 contributed three data points for the remaining 10 dimensions. This combined data provided a comprehensive measure of moral emotion intensity across all dimensions (see **Figure S1**).

**Figure S2.**
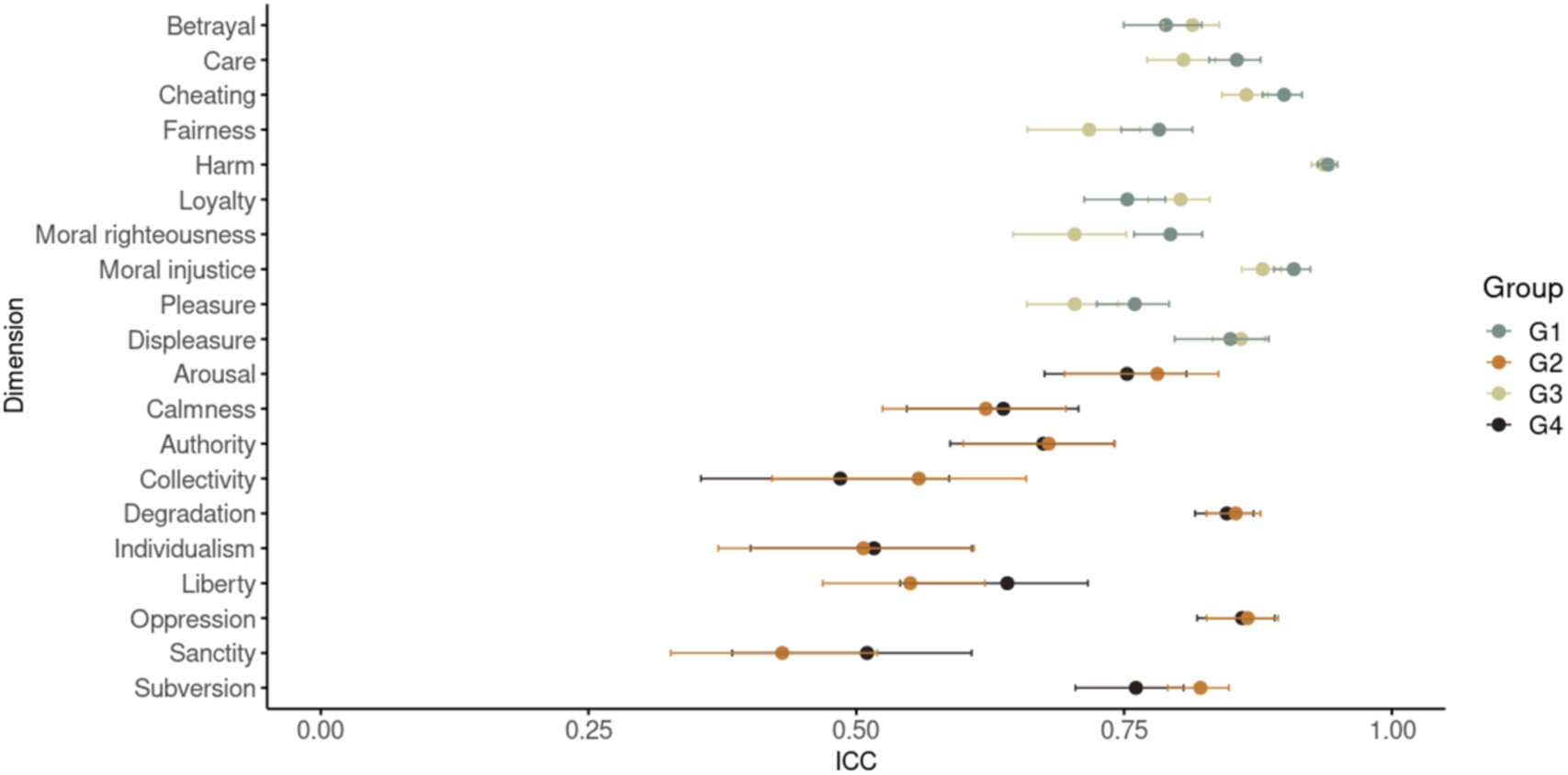
Intraclass correlation coefficients (ICC) for each rating group across all dimensions. Each point represents the ICC value, with the line on the left indicating the lower confidence interval and the line on the right indicating the upper confidence interval. The ICC was calculated using the R package psych. ICC(2k) was selected as the appropriate model because all raters within each group evaluated all video clips.

**Figure S3.**
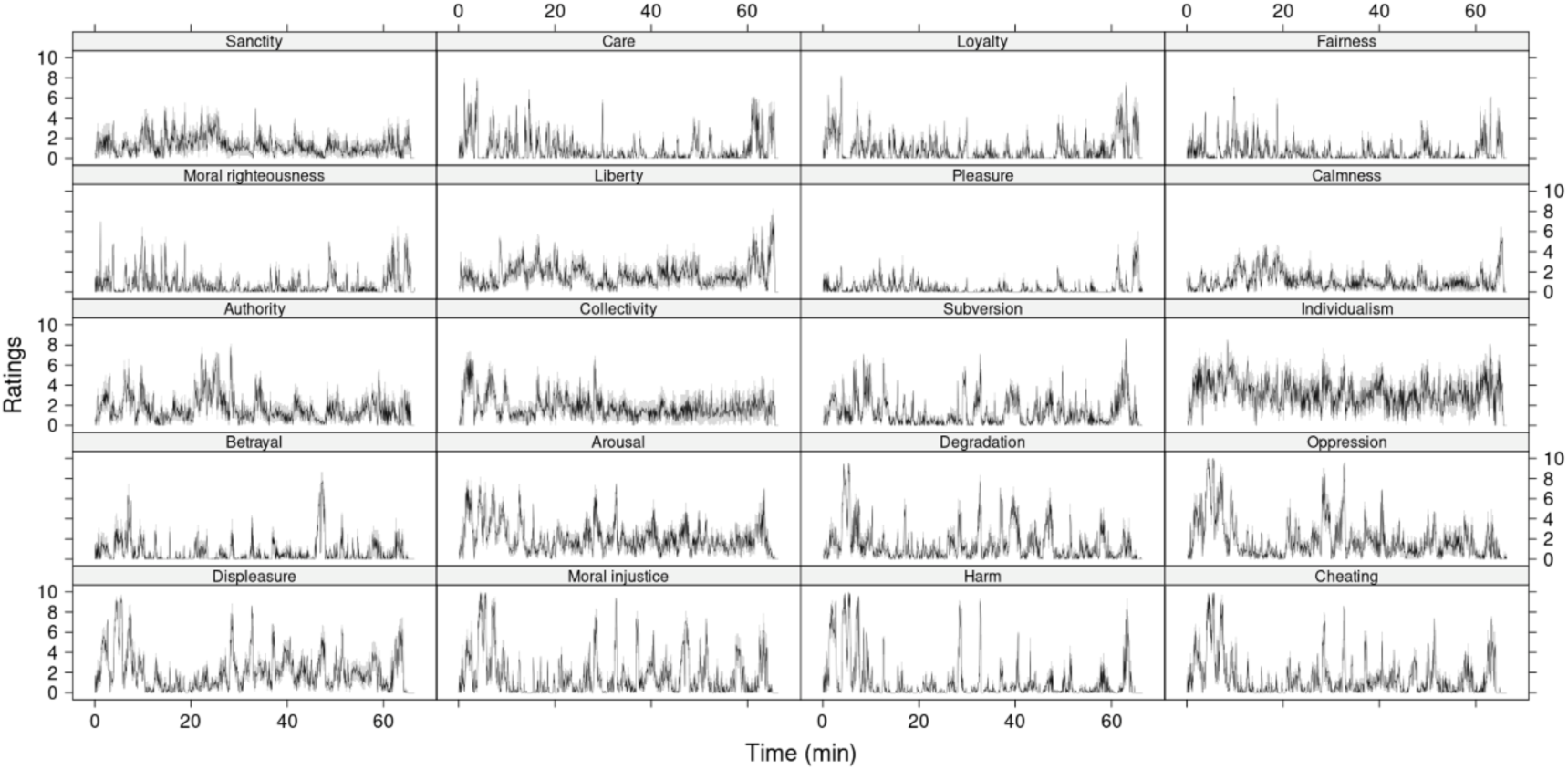
Moral inference ratings for 20 dimensions. The black line represents the mean ratings across participants, while the gray area indicates the standard error.

### Correlations and VIF values for the movie

**Table S1** presents the correlation matrices for each cluster during the first and second halves of the movie. The upper-right section of the table corresponds to the first half, while the lower-left section represents the second half.

**Table S1.** Correlation matrix of the clusters across the movie.

|  | Virtue | Hierarchy | Rebellion | Vice |
| --- | --- | --- | --- | --- |
| Virtue | 1 | 0.2385 | 0.1650 | -0.3428 |
| Hierarchy | 0.3608 | 1 | 0.2628 | 0.3552 |
| Rebellion | 0.4695 | 0.1873 | 1 | 0.5374 |
| Vice | -0.3189 | -0.0506 | 0.3917 | 1 |

**Table S2** provides the variance inflation factor (VIF) values for all clusters across the first and second halves of the movie.

**Table S2.** VIF values of the clusters across the movie.

|  | Virtue | Hierarchy | Rebellion | Vice |
| --- | --- | --- | --- | --- |
| First | 1.6971 | 1.3926 | 1.7663 | 2.3889 |
| Second | 2.2783 | 1.1571 | 2.2091 | 1.9265 |

### Inter-subject correlation analysis

Viewing movies featuring social interactions has been shown to synchronize brain activity across individuals (Hasson et al., 2004). However, how neural synchronization relates with moral inference remains unclear. To investigate this, we calculated the inter-subject correlation (ISC) of neural activation patterns using the ISC-toolbox (Kauppi et al., 2014). Dynamic ISC was calculated in consecutive time windows (window length: 10 TR = 26 seconds) to allow correlating time-window ISC with moral evaluations of the movie stimuli.

### Neural synchronization during moral inference

Whole-brain ISC analysis revealed extensive neural synchronization across participants during movie viewing (**Figure S4 & S5**).

**Figure S4.**
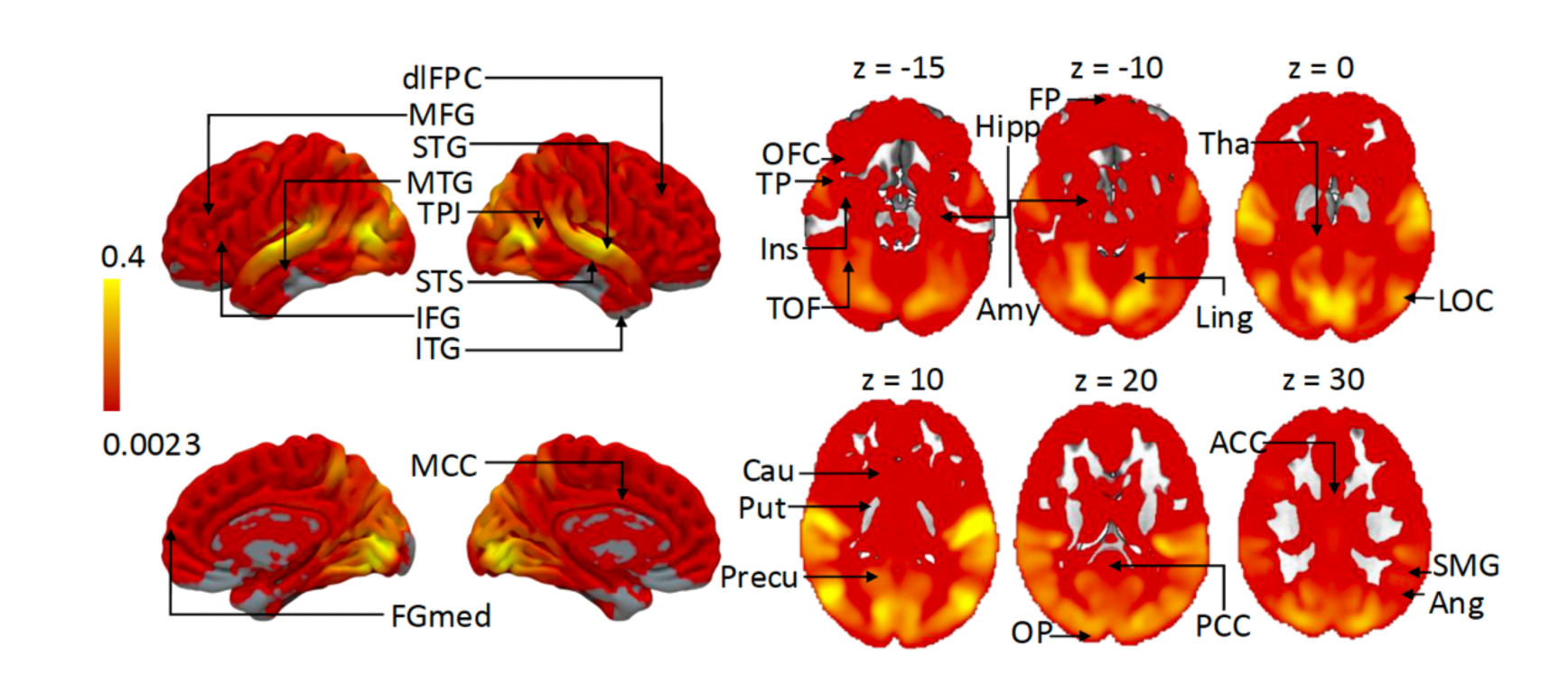
Significant ISC (FDR-corrected, *q* = 0.001) across subjects over the first half of the movie. dlPFC = dorsolateral prefrontal cortex, MFG = middle frontal gyrus, STG = superior temporal gyrus, MTG = middle temporal gyrus, TPJ = temporoparietal junction, STS = superior temporal sulcus, IFG = inferior frontal gyrus, ITG = inferior temporal gyrus, FGmed = medial frontal gyrus, MCC = middle cingulate cortex, OFC = orbitofrontal cortex, TP = temporal pole, Ins = insula, TOF = temporal occipital fusiform cortex, Hipp = hippocampus, FP = frontal pole, Amy = amygdala, Ling = lingual gyrus, Tha = thalamus, LOC = lateral occipital cortex, Cau = caudate, Put = putamen, Precu = precuneus, OP = occipital pole, PCC = posterior cingulate cortex, ACC = anterior cingulate cortex, SMG = supramarginal gyrus, Ang = angular gyrus.

**Figure S5.**
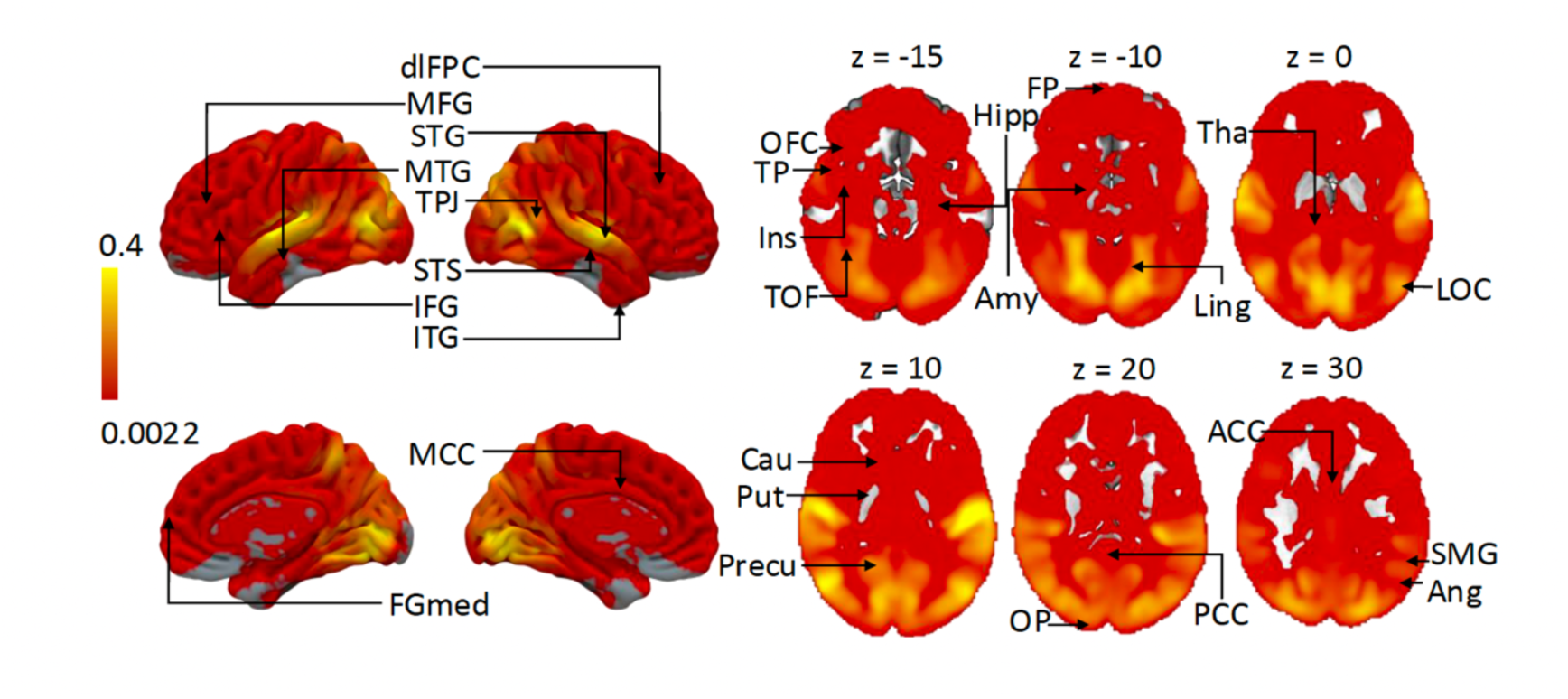
Significant ISC (FDR-corrected, *q* = 0.001) across subjects over the second half of the movie. dlPFC = dorsolateral prefrontal cortex, MFG = middle frontal gyrus, = superior temporal gyrus, MTG = middle temporal gyrus, TPJ = temporoparietal junction, STS = superior temporal sulcus, IFG = inferior frontal gyrus, ITG = inferior temporal gyrus, FGmed = medial frontal gyrus, MCC = middle cingulate cortex, OFC = orbitofrontal cortex, TP = temporal pole, Ins = insula, TOF = temporal occipital fusiform cortex, Hipp = hippocampus, FP = frontal pole, Amy = amygdala, Ling = lingual gyrus, Tha = thalamus, LOC = lateral occipital cortex, Cau = caudate, Put = putamen, Precu = precuneus, OP = occipital pole, PCC = posterior cingulate cortex, ACC = anterior cingulate cortex, SMG = supramarginal gyrus, Ang = angular gyrus.

**Figure S6** illustrates the brain regions where synchronization among participants correlates with the Vice and Rebellion main dimensions during the movie.

**Figure S6.**
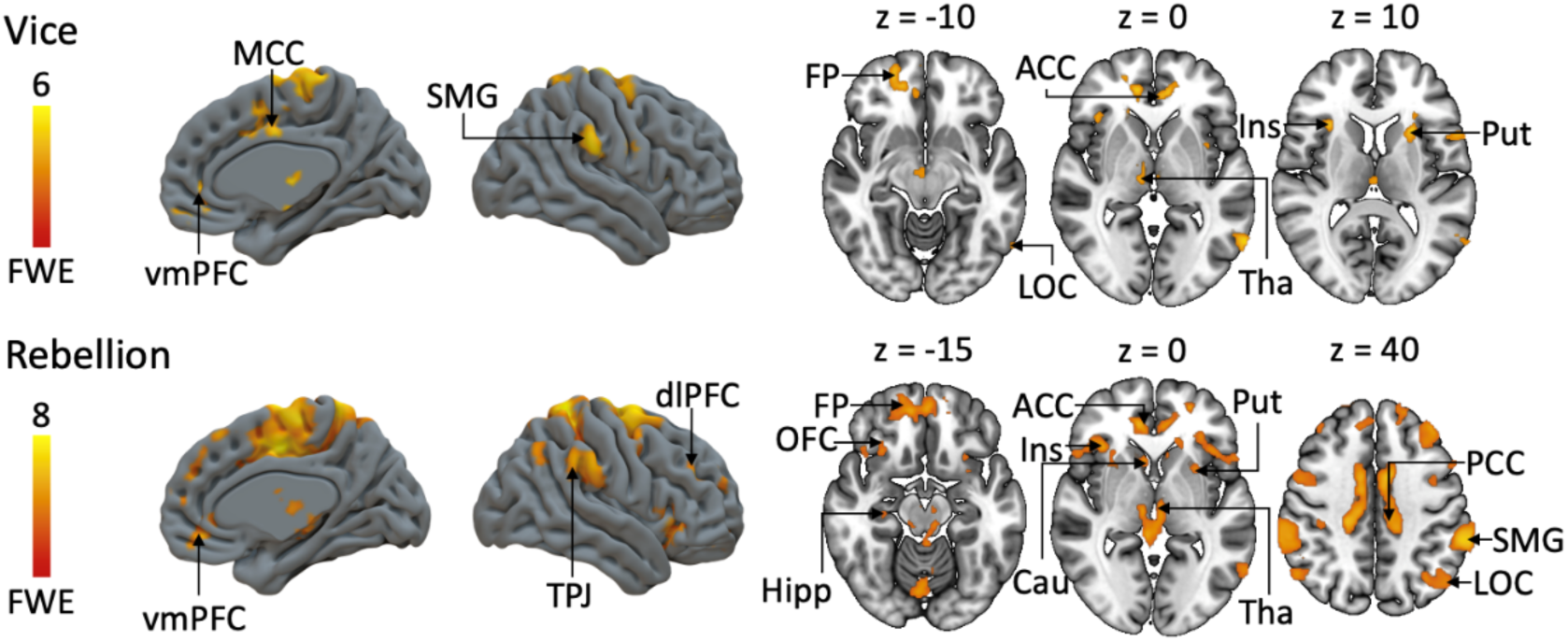
The significant associations between dynamic ISC and moral clusters (Cluster FWE-corrected at p < 0.05. Initial cluster-forming threshold p < 0.001). MCC = middle cingulate cortex, vmPFC = ventral medial prefrontal cortex, SMG = supramarginal gyrus, FP = frontal pole, LOC = lateral occipital cortex, ACC = anterior cingulate cortex, Tha = thalamus, Ins = insula, Put = putamen, dlPFC = dorsolateral prefrontal cortex, TPJ = temporoparietal junction, OFC = orbitofrontal cortex, Hipp = hippocampus, Cau = caudate, PCC = posterior cingulate cortex.

**Table S3.** Brain regions presented in Figure 7.

| Abbreviation | Full name |
| --- | --- |
| PCG | Precentral Gyrus |
| SFG | Superior Frontal Gyrus |
| MFG | Middle Frontal Gyrus |
| IFG-Op | Inferior Frontal Gyrus (Opercular Part) |
| IFG-Tri | Inferior Frontal Gyrus (Triangular Part) |
| IFG-Orb | Inferior Frontal Gyrus (Orbital Part) |
| ROL | Rolandic Operculum |
| SMA | Supplementary Motor Area |
| OLF | Olfactory Cortex |
| SFM | Superior Medial Frontal Cortex |
| MOF | Middle Orbital Frontal Cortex |
| REC | Rectus Gyrus |
| mOFC | Medial Orbitofrontal Cortex |
| aOFC | Anterior Orbitofrontal Cortex |
| pOFC | Posterior Orbitofrontal Cortex |
| lOFC | Lateral Orbitofrontal Cortex |
| INS | Insular Cortex |
| ACC | Anterior Cingulate Cortex |
| MCC | Middle Cingulate Cortex |
| PCC | Posterior Cingulate Cortex |
| HIP | Hippocampus |
| PHG | Parahippocampal Gyrus |
| AMY | Amygdala |
| CAL | Calcarine |
| CUN | Cuneus |
| LING | Lingual Gyrus |
| SOG | Superior Occipital Gyrus |
| MOG | Middle Occipital Gyrus |
| IOG | Inferior Occipital Gyrus |
| FFG | Fusiform Gyrus |
| PoCG | Postcentral Gyrus |
| SPL | Superior Parietal Lobule |
| IPL | Inferior Parietal Lobule |
| SMG | Supramarginal Gyrus |
| ANG | Angular Gyrus |
| PCUN | Precuneus |
| PCL | Paracentral Lobule |
| CAU | Caudate Nucleus |
| PUT | Putamen |
| PAL | Pallidum |
| THA | Thalamus |
| HES | Heschl |
| STG | Superior Temporal Gyrus |
| TPOsup | Temporal Pole (Superior) |
| MTG | Middle Temporal Gyrus |
| TPOmid | Temporal Pole (Middle) |
| ITG | Inferior Temporal Gyrus |

